# Discriminative histone imputation using chromatin accessibility

**DOI:** 10.1101/2024.01.11.575175

**Authors:** Wen Wen, Jiaxin Zhong, Zhaoxi Zhang, Lijuan Jia, Tinyi Chu, Nating Wang, Charles G. Danko, Zhong Wang

## Abstract

Histone modifications (HMs) play a pivot role in various biological processes, including transcription, replication and DNA repair, significantly impacting chromatin structure. These modifications underpin the molecular mechanisms of cell-specific gene expression and complex diseases. However, annotating HMs across different cell types solely using experimental approaches is impractical due to cost and time constraints. Herein, we present dHICA (discriminative histone imputation using chromatin accessibility), a novel deep learning framework that integrates DNA sequences and chromatin accessibility data to predict multiple HM tracks. Employing the Transformer architecture alongside dilated convolutions, dHICA boasts an extensive receptive field and captures more cell-type-specific information. dHICA not only outperforms state-of-the-art baselines but also achieves superior performance in cell-specific loci and gene elements, aligning with biological expectations. Furthermore, dHICA’s imputations hold significant potential for downstream applications, including chromatin state segmentation and elucidating the functional implications of SNPs. In conclusion, dHICA serves as an invaluable tool for advancing the understanding of chromatin dynamics, offering enhanced predictive capabilities and interpretability.

## Introduction

At the core of chromatin architecture are the highly conserved histone proteins—H1, H2A, H2B, H3, and H4—which serve as fundamental building blocks for packaging eukaryotic DNA into repetitive nucleosomal units[1]. These units are subsequently folded into higher-order chromatin fibers[2]. Histone modifications (HMs) significantly influence a broad range of cellular processes, including gene expression, chromatin structure modulation, and DNA repair[3, 4]. To elucidate the genome-wide signals associated with different cell types and tissues, initiatives like the Encyclopedia of DNA Elements(ENCODE)[5, 6] and the Roadmap Epigenomics Consortiums[7] have made substantial strides in systematically characterizing in vivo biochemical signatures across different cell types and tissues, including histone modifications, chromatin accessibility, and DNA methylation.

Despite these efforts to comprehensively map the epigenomes, significant challenges remain. Due to the high costs and time-consuming nature of experimental work, data have only been collected for a fraction of potential cell type and assay combinations outlined in these projects. Furthermore, considering the myriad developmental stages and environmental conditions, the diversity of possible human cell types is virtually boundless. It is impractical to anticipate collecting exhaustive data for every potential cell type/assay combination. Furthermore, no high-throughput assay is perfectly reproducible, and run-to-run differences in the same experiment may reflect either biological variation in the cells being assayed or experimental variance arising from sample preparation or downstream steps in the protocol.

As a practical solution, the development of in silico models to impute unknown epigenomic profiles based on existing data offers a promising alternative to these experimental limitations. Epigenomic imputation methods such as ChromImpute[8] and PREDICTD[9] have been presented to use available data to accurately impute the outcomes of missing experiments, thereby extending our understanding of epigenomic regulation across a more comprehensive spectrum of cell types and conditions. Alongside imputing missing data, Avocado[10] has produced a dense and information-rich representation of the human epigenomes, reducing redundancy, noise, and bias.

Considering the extensive range of tissues and histone modifications, relying solely on biological experiments to explore underlying mechanisms is somewhat unrealistic. Certain studies have been conducted to identify HM peaks, helping researchers to focus on regions more closely associated with regulatory effects on gene expression. For instance, DeepSEA[11] models regulatory information encoded by the DNA sequence to predict a wide array of epigenomics data, including TF-binding, DNase I sensitivity, and HM sites. Both DeepHistone[12] and iHMnBS[13] combine DNA sequence and DNase-seq data to classify multiple HM peaks. Meanwhile, DeepPTM[14] uses TF-binding data and DNA sequences to predict histone post-translational modifications; however, it is limited to predicting only a single modification marker in the center of the sequence for a given cell line. Yet, these methods are trained only in regions with HM peaks, not genome-wide, which may prevent them from capturing features across the entire genome.

Several models have been developed to predict genome-wide, cell-type-specific epigenetic and transcriptional profiles in large mammalian genomes. For instance, Kelley developed Basenji2[15], a deep learning model that predicts experimental HMs using DNA sequence alone. Enformer[16] advanced this approach by integrating a transformer architecture into the convolutional blocks, allowing it to process a larger range(197-kbp) of DNA sequences through the self-attention mechanism, handling longer sequences than Basenji2(131-kbp).

However, while DNA sequence encodes regulatory information for cells and tissue types, Enformer falls short in capturing the highly cell-type and developmental stage-specific nature of HMs[17]. dHIT[18] has shown promising outcomes in predicting HMs from GRO-seq data, demonstrating the potential of leveraging single-assay, cell-type-specific features to enhance predictive accuracy. EPCOT[19] incorporates DNase-seq and DNA sequence to predict HM tracks for a given cell type, utilizing a pre-training and fine-tuning framework. However, the resolution of EPCOT (1kbp) is significantly lower than that of Enformer and Basenji2, both of which are 128-bp, far underperforming the popular chromatin segmentation method (ChromHMM[20]) which uses 200-bp resolution. It should be noted that achieving higher resolution is crucial for various downstream applications, including chromatin segmentation.

Motivated by these insights, we introduce discriminative Histone Imputation using Chromatin Accessibility (dHICA) to predict multiple histone modification levels simultaneously using both DNA sequence data and chromatin accessibility as inputs. More importantly, the incorporation of the Transformer architecture [21] into our model expands its receptive field, allowing it to capture distal information. In cross-cell line and species evaluations, dHICA outperformed other state-of-the-art methods, primarily due to its effective integration of chromatin accessibility data. We discovered that chromatin accessibility data are crucial for model predictions, particularly for active marks. Unlike models that depend solely on DNA, dHICA can predict HMs in new cell lines and species without the need for re-training. Furthermore, dHICA’s imputed data can be utilized for downstream applications, including segmenting chromatin states and distinguishing histone acetylation quantitative trait loci (haQTLs) from SNPs. dHICA’s robust performance and versatility highlight its potential as an innovative tool in genomic research.

## Materials and methods

### Dataset

In this study, multiple HMs and sequencing files from ATAC-seq and DNase-seq were sourced from ENCODE[22]. The HMs utilized, based on the human genome version GRCh37, included H3K122ac, H3K4me1, H3K4me2, H3K4me3, H3K27ac, H3K27me3, H3K36me3, H3K9ac, H3K9me3, and H4K20me1. These markers are indicative of specific functional elements such as enhancers, promoters, and gene bodies. This study also explored less commonly studied modifications like H3 lysine 122 acetylation (H3K122ac). For the experimental setup, the K562 cell line (chromosomes 1-21) was used for model training and validation. Chromosome 22 of the K562 line, along with other cell lines such as GM12878 and HCT116, served as the testing ground. Two separate models were optimally trained using DNase-seq and ATAC-seq data respectively. Tables S2 and S3 in the supplementary material provide further details on the data used.

### Data preprocessing

In this study, mitochondrial DNA is excluded from the analysis, focusing only on the autosomes and sex chromosomes. To minimize potential confounding factors, ENCODE blacklist regions are also omitted from the entire genome[23]. Avoiding assembly gaps and unmappable regions greater than 1kb, we extract 131-kb non-overlapping sequences across the chromosomes, which expand to 197-kb on both sides as model inputs. This procedure yielded 22,727 intervals for extracting DNA sequences and chromatin accessibility data.

DNA sequences are read using a one-hot encoding scheme, where A = [1,0,0,0], C = [0,1,0,0], G = [0,0,1,0], T = [0,0,0,1], and N = [0,0,0,0]. Chromatin accessibility data is extracted directly from fold change bigwig files, and we ensure data integrity by setting any negative or NaN values to zero, without applying any further data transformation. For predicting HM signals, within each interval, we summed coverage estimates in a bin with the length of 128-bp to serve as the signal for the model to predict.

To mitigate the influence of experimental factors such as batch effects and data quality variations, four distinct chromatin accessibility datasets are utilized for the same set of interval partitions. This method not only captures the variability across different experimental conditions but also effectively quadruples the sample size, thereby enhancing the robustness of the data processing. Consequently, the assembled dataset encompasses a comprehensive total of 91,908 samples(exactly fourfold the base count of 22,727 intervals) with 87,868 samples designated for training, 1,472 for validation, and 1,568 for testing, thereby guaranteeing extensive coverage and reliable model evaluation.

### Architecture of the dHICA

Our proposed model, dHICA, builds upon the foundation of Enformer, and integrates chromatin accessibility data with DNA sequence to enhance the prediction of HMs, as illustrated in Figure 1A. The architecture comprises a sophisticated sequence of layers designed for optimal data processing and prediction accuracy. It starts with two separate convolutional blocks, one dedicated to DNA sequences and the other to chromatin accessibility data. Each block is tailored to extract the specific characteristics of its input data type. These features are then processed through a fusion layer, which prepares them for the next critical phase. The Transformer block, a pivotal component of the model, excels in capturing long-range dependencies and interactions between the DNA sequences and chromatin accessibility, crucial for understanding their combined influence on HMs. The processing sequence concludes with a cropping layer followed by a fully connected layer, which together refine and predict the 10 specific types of HMs. This integrated approach ensures that dHICA not only captures the unique aspects of each data but also effectively interprets the complex interdependencies that dictate histone modification patterns.

**Fig. 1.**
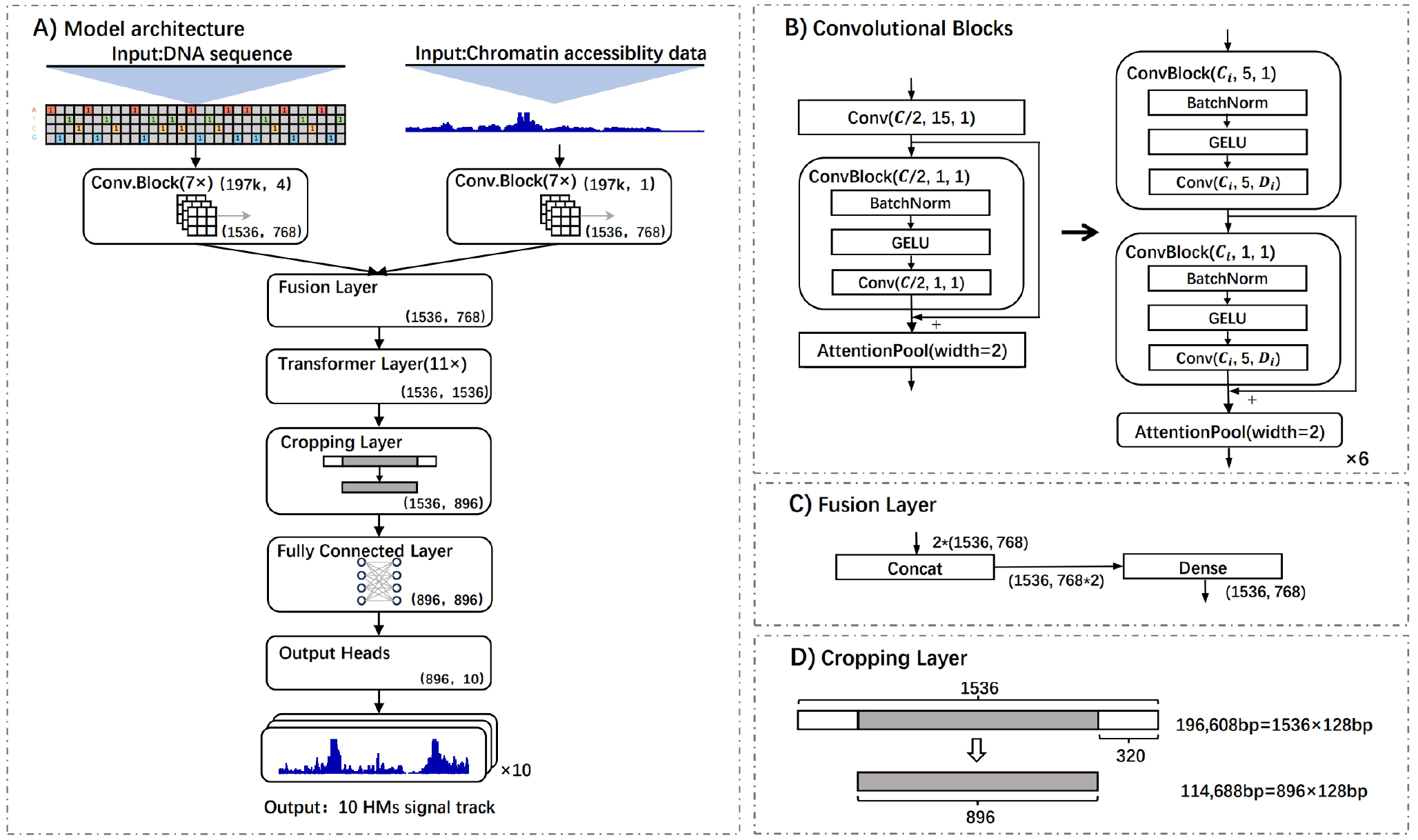
The dHICA Framework Illustrated. **(A)** Overview of the dHICA. **(B)** Detailed depiction of the convolutional blocks within dHICA. **(C)** The fusion layer of dHICA, integrating features derived from both DNA sequence and chromatin accessibility data. **(D)** The strategy dHICA employs to segment distant genomic regions.

### Convolutional blocks and fusion layer

We applied convolutional blocks with pooling to distill the input data, DNA and chromatin accessibility data, into fixed-size representations. Specifically, the model dissects the input sequences into 128-base pair(bp) bins to achieve the desired resolution.

The architecture comprises seven distinct parts within the convolutional blocks, these are categorized into two primary convolutional blocks as depicted in Figure 1B. The initial convolutional block effectively condenses the spatial dimension from 196608-bp down to 1536, which ensures that each vector in the sequence symbolizes a 128-bp bin aligning with the resolution parameters set for dHICA.

Following this, six additional blocks utilize dilated convolutions–a technique where the convolutional filters incorporate deliberate gaps progressively enlarged by a factor of two in each subsequent layer. This approach allows the model’s receptive field to expand exponentially without increasing the complexity linearly. A key feature of this model is the dense connectivity of these layers, whereby each layer utilizes inputs from all preceding layers instead of only the previous one. This design optimizes the number of filters required per layer as it allows for the preservation and integration of the rich feature set extracted from the initial convolutional operations through to the complex nuances teased out by the dilated convolutions. Each layer can thus concentrate on capturing the residual variation that previous layers have not addressed.

For the dilated convolution layers, we increase the number of channels *C*_*i*_ by a consistent multiplier until we attain the specified channel count *C*, starting from half that value *C/*2 in the initial six layers. In tandem, the dilation rate *D*_*i*_ is augmented by a factor of 1.5 for each successive layer, with the resulting figure rounded to the nearest whole number. For our setup, we define the channel size *C* as 1536 and initiate with *C/*2 filters along with a pooling size of 2.

To further refine and integrate the features extracted from the DNA and chromatin accessibility data, we employ fusion layers(Figure 1C). The first dense layer serves to double the channel capacity, thereby amplifying the feature space, while the subsequent layer scales it back to the model’s baseline number of channels. This methodical expansion and contraction of the channel space facilitate a more nuanced synthesis of the underlying biological signals.

### Transformer block

The Transformer block is the core component of the model encompassing 11 distinct layers that each play a pivotal role in interpreting sequence data. This block operates by transforming each position in the input sequence through a computed weighted sum of all position representations, a process known as attention. Here, attention weights are influenced by the embeddings of the positions and their relative distances, enabling the incorporation of spatial context.

This attention-driven mechanism is key to the model’s advanced ability to predict HMs. It leverages information from critical regulatory regions such as enhancers, which are essential for gene regulation. A standout feature of the model is its ability to focus attention directly across the entire sequence, facilitating seamless information exchange across potentially distant elements along the DNA strand. As a result, the Transformer layers significantly broaden the model’s receptive field, capturing regulatory elements up to 100 kb away, while preserving the integrity of information crucial for accurate predictions.

The attention mechanism within these blocks is mathematically represented as:

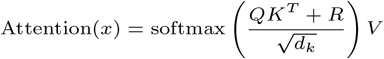

where *Q, K, V* represent the queries, keys, and values vectors—each a critical component of the attention calculation; *R* denotes the relative positional encodings, integrating spatial context into the model; and *d*_*k*_ is the dimensionality of the keys and queries, providing necessary scaling. The softmax function ensures that the attention weights are normalized across the sequence.

Furthermore, the model employs Multi-Head Attention (MHA) to conduct multiple attention computations in parallel, allowing each head to independently capture distinct features of the input data:

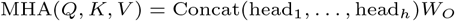

where each head_*i*_ = Attention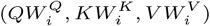 and 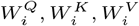 are parameter matrices for *i*-th attention head, *W*_*O*_ is the output weight matrix that combines the heads; the number of heads *h* is set to 8.

### Cropping layer

To tackle the computational challenges associated with analyzing distant regions in genomic sequence data, a cropping layer is employed. This layer excises 320 positions from each end of the sequence, effectively shortening it by 320 *×* 128 bp as illustrated in Figure 1D. The process leaves only the central 896 positions. This cropping strategy is crucial because of the model’s inherent limitations in capturing and learning effectively from regions distant from the sequence center. These constraints are due to the model’s architecture, which is designed to primarily perceive and analyze regulatory elements facing towards the sequence center, while it struggles to detect elements beyond the sequence boundaries.

The cropping operation is mathematically depicted as follows:

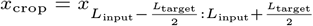

where *x* is the output from Transformer blocks, *L*_input_ is the original sequence length; *L*_target_ is the target sequence length after cropping.

Following this, a fully connected layer is implemented to predict HMs with a resolution of 128-bp. This step not only refines the model’s output but also significantly reduces the computational burden.

### Model training

We adopted a supervised learning approach to optimize our dHICA model, utilizing the Mean Squared Error (MSE) loss function to optimize performance. To assess which chromatin accessibility data better complements dHICA, we conducted parallel training sessions using identical model architectures on both ATAC-seq and DNase-seq data. Our training dataset comprised chromosomes 1-20 from the K562 cell line, with chromosome 21 reserved for validation and chromosome 22 for testing.

We configured the MSE loss function with an initial learning rate of 0.0001, incorporating a learning rate decay strategy that reduces the rate by a factor of 1.4 every 5 epochs following the first 10 epochs. The model was optimized over 300 epochs, processing batches of 1,500 samples each. This rigorous training regimen ensures that dHICA can reliably predict steady-state HMs across various cell types, assuming the underlying relationships between HMs, DNA, and chromatin accessibility signals remain consistent.

## Results

### Performance evaluation across different cell lines, tissues and species

To thoroughly investigate the generalization capability of our model, we applied dHICA on a diverse array of cell lines(GM12878, MCF-7, HeLa-S3, HCT116, HepG2, IMR-90 and A549), along with human(Heart and Spleen) and mouse (Hindbrain, Heart and G1E) tissues, despite its exclusive training on K562 cell line.

The imputed HMs via dHICA exhibit robust correlation with experimental data (Figure 2A)[24]. Particularly noteworthy are the disparities between the imputed and experimental signals, frequently observed in background regions with low signal intensity, suggesting the presence of unaccounted technical variations in the ChIP-seq background signal[25], distinct from chromatin accessibility data. Among the marks we attempted to model, only the repressive marks H3K9me3 and H3K27me3 showed subpar performance, likely due to their weak correlation with chromatin accessibility signals, low data values, and average sequencing quality[26, 18].

**Fig. 2.**
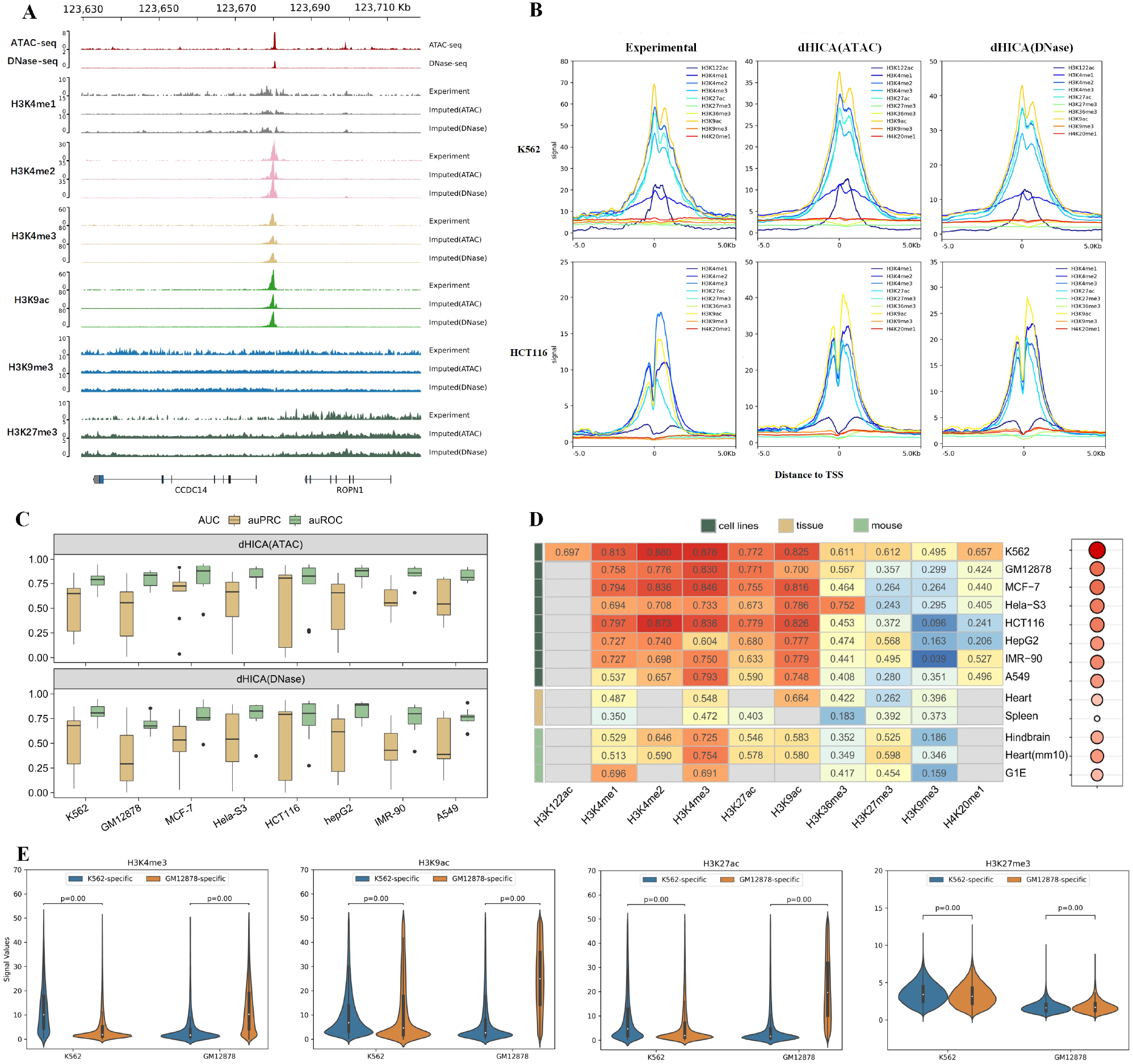
Comprehensive analysis of cross-cell-type and cross-species imputations. **(A)** Genome browser comparison between experimental and predicted histone marks near gene CCDC14 and ROPN1 in GM12878. **(B)** Comparison of dHICA’s imputed signals and experimental data proximal to the Transcription Start Site (TSS) in K562 and HCT116 cell lines. **(C)** Evaluation of dHICA’s performance across HM peak regions. **(D)** Comparison of Pearson’s correlation of ATAC model across cell lines, tissues and species. Empty cells indicate that no experimental data are available for comparison in the cell type shown. **(E)**Distribution of cell-type-specific HM imputations by dHICA, indicating the effective capability of dHICA in distinguishing cell-specific feature.

For the evaluation of HM imputations, we employed the method outlined in the ENCODE imputation challenge[27] to compute the Pearson’s correlation between imputed and experimental HMs across 7 distinct cell lines (Figure 2D, Figure S2). Active marks (H3K4me3, H3K4me2, H3K4me1 and H3K9ac), which are predominantly associated with promoters and enhancers, exhibited consistent performance across holdout cell types, akin to the performance observed in the training cell line K562(with an average Pearson’s correlation exceeding 0.7). However, there was a slightly decrease in performance near repressive regions(H3K9me3, H3K27me3 and H3K20me1). Moreover, the imputation performance across different cell lines correlated with the similarity of HMs in correlation between the predicted cell line and the training cell line K562 (Figure S1). Nevertheless, the model showcases robust generalization across all HMs, demonstrating its effectiveness in diverse chromatin contexts.

Moving beyond genome-wide predictive accuracy, we delved into the imputed results near HM peaks and transcription start sites(TSS). The signals in those regions are the most informative[28], enabling a detailed examination of their distribution. Notably, no significant disparities in accuracy were observed across various regions of HM peaks with high signal intensity in either experimental or imputed data(Figure 2C). Moreover, the imputation effectively captured the nuanced distribution of HM signals near the TSS of annotated genes (Figure 2B). Within TSS regions, dHICA comprehensively encompasses both active marks and repressive marks, aligning closely with biological expectations[29]. The substantial correlation between the distribution of imputed and experimental HM signals further bolsters confidence in dHICA’s ability to accurately represent biological phenomena, affirming that it goes beyond merely learning average signal intensities of HMs[30].

Additionally, our model, originally trained on the human K562 cell line, extended to HM signals in mouse cell and tissue types(Figure 2D). What surprised us is that dHICA achieve even higher accuracy in mouse tissues compared to human tissues, which was due to data quality issues. Each exhibiting distinct levels of similarity with the training cell line without retraining the model, which could help explore the general feature of HMs that shared across mammalian cell types. This capability unveils significant potential applications in annotating genomes of less-explored mammalian species.

Inspired by CEMIG[31], we employed dHICA to discern cell-specific HM peak sites in K562 and GM12878 cells. For each HM, we delineated K562-specific, GM12878-specific and shared peaks(detailed in supplementary Text S2). The variation in loci between different cell types reflects distinct regulatory mechanisms and gene expression patterns[32]. We investigate the ability of dHICA in figuring those cell-specific features by comparing the distribution of signal values in different cell-specific peak regions, measured in raw counts imputed by dHICA(Figure 2E). Ideally, the imputation for the K562 cell line should show significantly higher signal values at K562-specific peaks than at GM12878-specific peaks, with a reciprocal pattern expected for GM12878 cell line imputations. A one-sided Wilcoxon rank sum test against the null hypothesis that the signal values in different cell-specific peaks are identical yielded a p-value significantly less than 0.05, supporting the conclusion that dHICA effectively distinguishes cell-specific features. Despite being trained solely on the K562 cell line, dHICA accurately identifies specific regions, even in GM12878 cells where it hasn’t been trained.

### Performance comparison with state-of-art methods

Given dHICA’s ability to accurately predict epigenomic features across cell lines, to further evaluation, we compare it’s performance with other baseline methods, including EPCOT, Enformer, deepPTM, deepSEA and ChromImpute. However, due to diverse nature of prediction tasks(binary models for classifying HM peaks vs. quantitative models for imputing signal tracks), making direct fair comparisons is challenging[33]. To address this issue, we divided the comparison into two main aspects: one focusing on performance in HM peak regions, and the other assessing genome-wide performance.

To compare the performance of state-of-art methods in HM peak regions and due to data imbalance, we calculate the auPRC of imputations rather than auROC from different baseline methods, as shown in the Table 1, Figure S3 and S4. Given that deepPTM only imputes H3K4me3, H3K9ac, H3K27ac, and H3K27me3, we select these four HM markers for comparison, as they correlate with both active and repressive regions in the genome. This selection ensures the subsets of HMs are reasonable. Notably, dHICA using ATAC-seq achieved the best performance, with an average auPRC higher than 0.7. For genome-wide HM imputation comparison, we compute Pearson’s correlation, Spearman’s correlation and RMSE between experimental signals and imputation from dHICA, EPCOT and Enformer[26]. From the Figure 3A, we easily discern that dHICA outperforms other baselines in all performance metrics. We further compare dHICA with EPCOT, as it’s the most similar and comparable to our model. Like dHICA, it incorporates DNA sequences and chromatin accessibility data for predictive task. However, EPCOT used four cell lines (K562, MCF-7, GM12878 and HepG2) as training dataset, whereas dHICA only used K562 cell line. To make the comparison fairer, we calculate the correlation of HMs between different cell lines(Figure S1), and we ultimately select five cell lines that are most similar to K562, along with K562 itself for evaluation between EPCOT and dHICA. Among the 6 cell lines(K562, GM12878, MCF-7, HCT116, HeLa-S3 and IMR-90) used for comparison, dHICA only used one cell line, K562, for training, while EPCOT used three cell line(K562, GM12878, MCF-7) for model training. Therefore, EPCOT could be expected to be more effective than dHICA. Nevertheless, we consistently outperform EPCOT in the prediction of HM markers(Figure 3B), regardless of the type of chromatin accessibility signals used by the model (ATAC-seq or DNase-seq).

**Table 1.** The comparison of auPRC bewteen different models across HM peak regions.

| Methods | H3K4me3 | H3K9ac | H3K27ac | H3K27me3 | Average | Cell line |
| --- | --- | --- | --- | --- | --- | --- |
| dHICA(ATAC) | 0.919 | 0.708 | 0.718 | 0.575 | 0.730 | K562 |
| dHICA(Dnase) | 0.871 | 0.649 | 0.647 | 0.360 | 0.632 | K562 |
| EPCOT(ATAC) | 0.887 | 0.528 | 0.482 | 0.258 | 0.539 | K562 |
| EPCOT(DNase) | 0.857 | 0.531 | 0.637 | 0.359 | 0.596 | K562 |
| Enformer | 0.865 | 0.578 | 0.530 | 0.160 | 0.533 | K562 |
| deepPTM | 0.903 | 0.719 | 0.554 | 0.240 | 0.604 | H1 |
| deepSEA | 0.737 | 0.563 | 0.534 | 0.440 | 0.569 | H1 |
| ChromImpute | 0.617 | 0.688 | 0.200 | 0.788 | 0.573 | H1 |

**Fig. 3.**
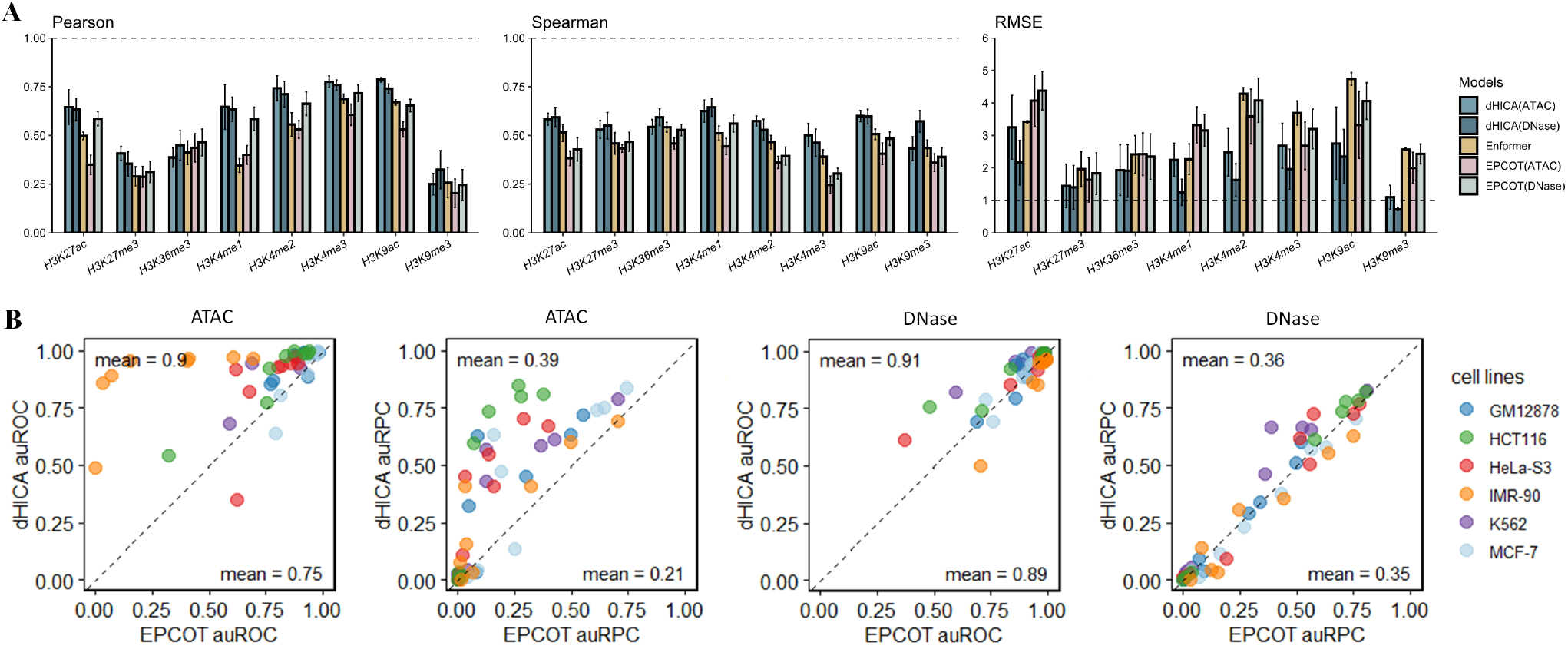
Multidimensional Performance Assessment. **(A)** Aggregate metrics including Pearson’s correlation, Spearman’s correlation, and Root Mean Square Error (RMSE) evaluated over six cell lines. **(B)** Genome-wide comparison of EPCOT and dHICA methodologies across multiple cell lines.

### Contribution of DNA and chromatin accessibility data

Most of the baselines rely solely on DNA sequence inputs, potentially lacking cell-specific features. In contrast, dHICA, integrates both one-hot encoded DNA sequences and cell-type specific chromatin accessibility signals. These cell-specific signals, represented by raw sequencing reads without any data transformation, are used to comprehensively impute HM signals. For cell-type specific inputs, we opt for ubiquitous chromatin accessibility profiles from DNase-seq and ATAC-seq due to their profound implications in gene regulation and chromatin organization.

To evaluate the impact of DNA information and chromatin accessibility data on enhancing the model’s performance in predicting HM signals, we separately trained the dHICA using only DNA or chromatin accessibility data on K562 cell line. Subsequently, we assessed the performance of these individual models on the HCT116 cell line and compared them with the standard dHICA model. Initially, we computed genome-wide Pearson’s correlation from imputations generated by different individual models. As depicted in Figure 4A, using both DNA and chromatin accessibility data consistently yielded higher Pearson’s correlation compared to using either component alone. Notably, for active marks, relying solely on chromatin accessibility data outperformed the use of DNA sequence data alone, whereas this was not the case for repressive marks.

**Fig. 4.**
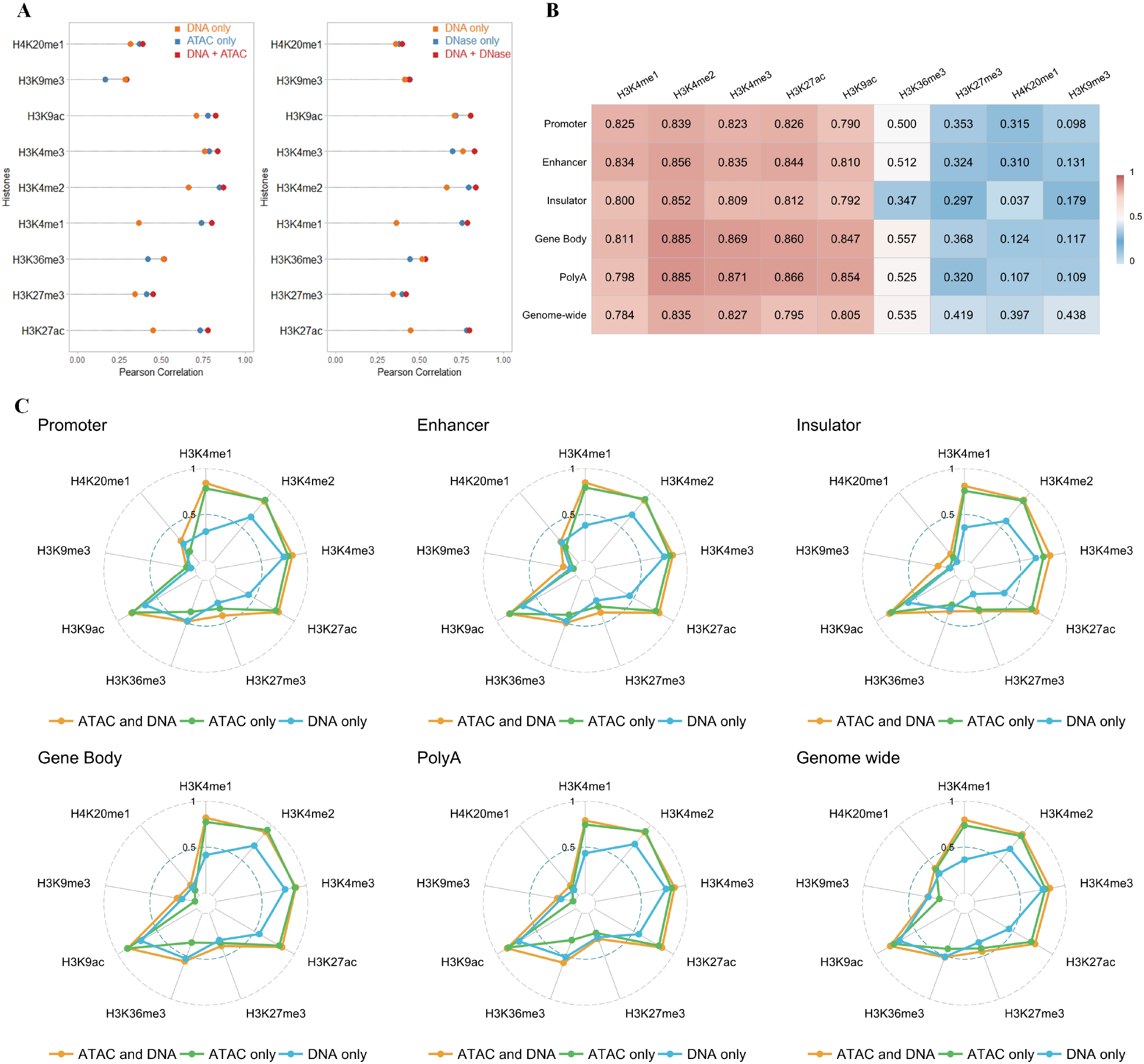
Analysis of the contribution of DNA and chromatin accessibility data. **(A)** Pearson’s correlation of dHICA genome-wide using DNA or chromatin accessibility data in HCT116 cell line. **(B)**The performance of dHCIA’s imputations across different gene elements in HCT116 cell line. **(C)** The contribution of DNA and chromatin accessibility data across different gene elements in HCT116 cell line.

Considering the pivotal role of gene elements and their intricate interactions with HMs[34], along with the use of HM signals near various gene elements by many computational methods to predict gene expression[35, 36, 37], we are particularly interested in the predictive performance of HMs around gene elements. We finally selected five gene elements: promoter, enhancer, insulator, gene body and PolyA, and calculate the Pearson’s correlation in these regions(Figure 4B). Aligned with the intricate interaction between HMs and gene elements, HMs exhibit significant performance particularly in regions where they closely associate with specific gene elements. For instance, H3K4me3 in promoters, H3K4me1 in enhancers, and H3K36me3 in gene bodies.

To delve deeper into the primary impact of DNA information and chromatin accessibility data in gene elements, we conducted a comparative analysis of the Pearson’s correlation of the individual models around these gene elements, as depicted in Figure 4C. Consistent with the genome-wide conclusion, for markers associated with enhancers and promoters, chromatin accessibility data plays a more crucial role than DNA. Conversely, for marks associated with transcription and repressive regions, DNA assumes greater importance. However, the incorporation of chromatin accessibility data also contributes significantly, as evidenced by the superior performance of the standard dHICA model compared to individual models using DNA alone genome-wide.

### dHICA enables precise chromatin state imputation for landscape insights

Chromatin state segmentation and genome annotations are essential for various genomic tasks, including the identification of active regulatory elements and the interpretation of disease-associated genetic variations across different cell types and in human disease[38, 39]. Given the robust performance of dHICA in the vicinity of gene elements, we investigated whether chromatin states defined by ChromHMM could be inferred using HM markers imputed by dHICA[40]. We used the pre-trained reported ChromHMM model that defined 18 distinct chromatin states based on six marks for which we trained imputation models(H3K4me3, H3K27ac, H3K4me1, H3K36me3, H3K9me3 and H3K27me3)[20]. Examination through the Integrative Genomics Viewer(IGV) showed that chromatin states were highly consistent[41], regardless of whether they were defined using ENCODE data or dHICA’s imputation(Figure 5B). This highlights dHICA’s ability to accurately represent the underlying epigenetic landscape, successfully extrapolating complex chromatin configurations from integrated datasets. The consistency of the imputed states with those derived from ENCODE data suggests that dHICA can be a viable alternative for predicting chromatin states, particularly in contexts where ChIP-seq data is unavailable or when detailed analysis of chromatin states is required due to the extensive variability in human chromatin states[42].

**Fig. 5.**
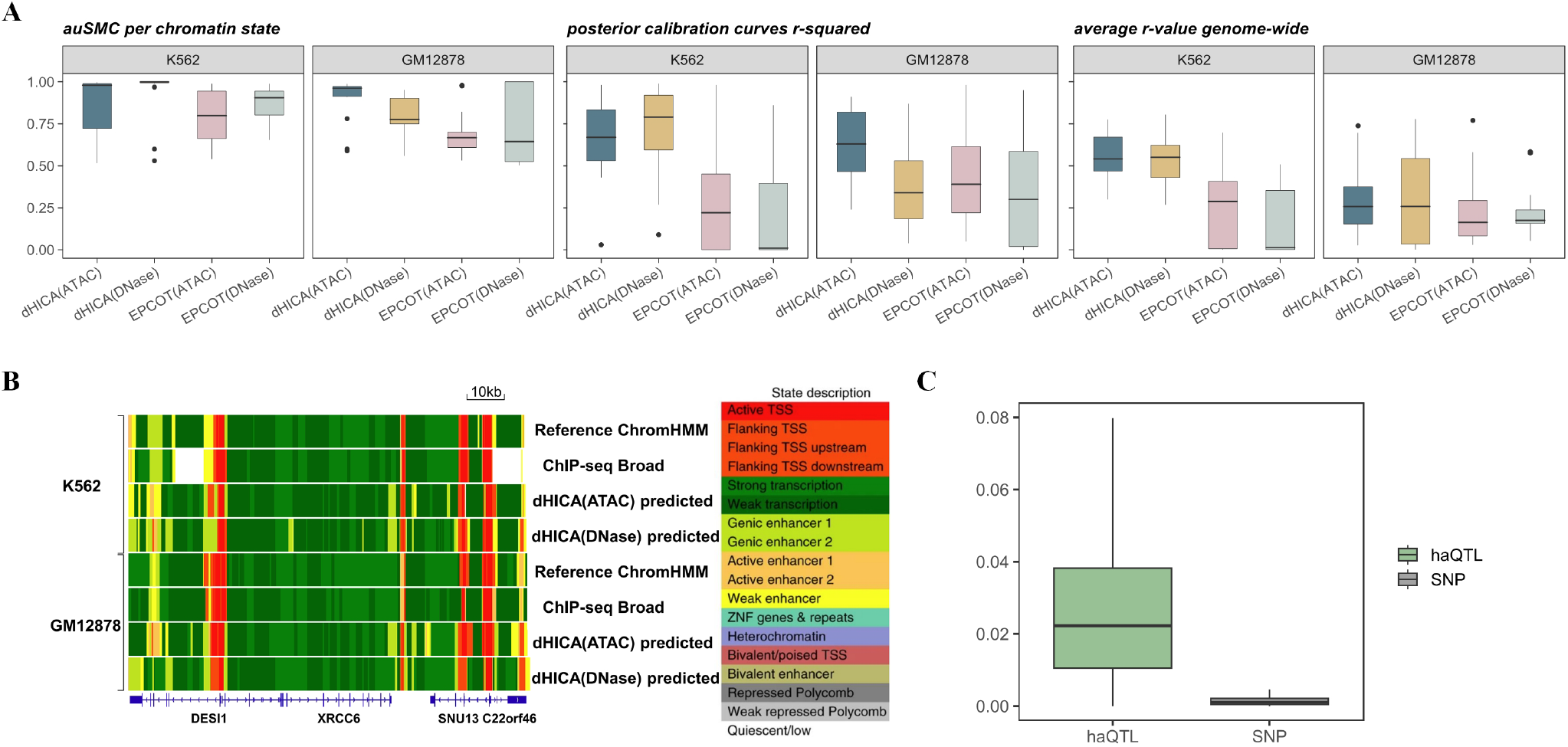
Downstream application and explanation of dHCIA. **(A)** Performance assessment for chromatin state segmentation using ChromHMM based on HM signals imputed by dHICA and EPCOT. **(B)**Genome browser in K562 and GM12878 cells shows the 18-state ChromHMM model using ChIP-seq data used to train the model (Broad) or based on imputation (dHICA predicted). **(C)** Functional implication scores (FIS) of haQTLs and nearby SNPs within 500bp.

To achieve a more quantitatively robust and principled evaluation of chromatin state segmentation, we applied SAGAconf[43] to compare the annotations derived from imputation with ENCODE ChIP-seq data in both K562 and GM12878 cell lines. We included EPCOT for comparison, as it closely resembles our model. Illustrated in Figure 5A, we calculated the area under the scaled min-max curve (auSMC), posterior calibration curves r-squared, and correlation coefficients (r-values) between the ENCODE data and the imputations generated by both dHICA and EPCOT, with detailed metrics provided in the Supplementary Text S3. For each of the metric, dHICA outperformed EPCOT, regardless of whether ATAC-seq or DNase-seq data were used. Additionally, although GM12878 serves as a test cell type for dHICA and a training cell type for EPCOT, where EPCOT should presumably perform better, dHICA still excelled over EPCOT in segmentation and genome annotations task.

### dHICA explains functional implications of SNPs

Genome-wide association studies (GWAS) have successfully identified numerous genetic variants associated with complex traits and diseases[44, 45, 46]. However, elucidating the biological mechanisms underlying these associations is challenging, as most SNPs are noncoding and their regulatory roles remain unclear[47, 48]. The genotype-independent signal correlation and imbalance (G-SCI) test[49], leveraging ChIP-seq assays on H3K27ac, has streamlined the identification of histone acetylation quantitative trait loci (haQTLs), thereby aiding in pinpointing causal variants within GWAS loci and advancing our understanding of their functional implications.

Inspired by the G-SCI method and studies on cell-type-specific haQTLs[50, 51, 50], we employed the dHICA, which can impute cell-specific and tissue-specific HM signals, to analyze the SNPs identified by G-SCI, which demonstrates dHICA’s potential to enhance understanding of the functional implications of these SNPs. Following DeepHistone[12], we identified a set of 6925 SNPs (haQTLs) specific to H3K27ac in the GM12878 cell line from the 1000 Genomes Project[52]. We also created a negative control set with an equivalent number of SNPs, each approximately 500-bp away from a corresponding haQTL. Using the formula Δ*p* = |*p*_ref_ *− p*_alt_|, where *p*_ref_ denotes the signal intensity associated with the reference allele, and *p*_alt_ represents the signal intensity of the alternative allele post-mutation, as defined in the study [53], we calculated functional implication scores for these SNPs. The results, displayed in Figure 5C, indicate that haQTLs have significantly higher scores than control SNPs. This was substantiated by a one-sided Wilcoxon rank sum test, which revealed a marked difference in the median scores between the two groups, with a p-value *<*0.05. Consequently, our analysis supports the biological understanding that haQTLs are more likely to impact the function of the lymphoblastoid epigenome and, in turn, influence phenotypic traits[54, 55]. This demonstrates that dHICA can effectively distinguish SNPs that are potentially responsible for specific phenotypes from their neighboring genetic variants.

## DISCUSSION

In this study, we presented dHICA, a deep learning framework that integrates chromatin accessibility information and DNA sequences to accurately predict cell-specific HM signals. By incorporating the transformer structure alongside dilated convolutions, dHICA significantly expands the model’s receptive field, enabling it to capture long-range interactions between genomic elements. dHICA outperformed other state-of-the-art methods across various cell lines and species, primarily due to its integration of chromatin accessibility data, which provides cell-specific features, particularly active gene elements. Moreover, dHICA’s imputed data can be utilized for downstream tasks such as chromatin state segmentation genome annotation, and distinguishing haQTLs from SNPs. Given its ability to predict HMs in new cell lines and species without re-training, dHICA holds significant potential for refined and personalized analysis, provided that users can supply the necessary chromatin accessibility data.

While dHICA demonstrates superior performance, there are opportunities for further refinement. (1) data normalization and quality improvement: currently, dHICA processes raw counts directly from bigWig files without any data transformation or normalization, which may introduce noise and variability. Implementing normalization strategies for GC%[56], employing denoising tools[57], or exploring data transformation techniques such as arcsinh-transformed epigenomic feature signals could enhance data quality. (2) expansion across cell lines: dHICA is presently trained exclusively on the K562 cell line. Incorporating sequencing data from multiple cell lines could enrich the model with diverse, cell-type-specific features, thereby enhancing its predictive accuracy. (3) prediction of gene expression: given the established link between HMs and gene expression[58], and considering dHICA’s capability in accurately imputing HMs, there is potential for the model to predict gene expression. Ideally, this capability would significantly broaden dHICA’s applications and contribute to a deeper understanding of biological mechanisms.

## Key Points

- This study proposes a deep learning framework, dHICA, which integrates chromatin accessibility information and DNA sequences to accurately predict cell-specific HM signals.
- dHICA largely benefit from the chromatin accessibility data.
- Across various cell lines, dHICA consistently outperforms other state-of-the-art methods.
- dHICA facilitates precise chromatin state segmentation, providing deeper insights into the genomic landscape.
- dHICA aids in elucidating the functional implications of SNPs.

## Code availability

The code for dHICA is available on GitHub at https://github.com/wzhy2000/dHICA. We also offer a cloud computing service at https://dreg.dnasequence.org/. Through this platform, users can upload their data to use GPU resources and obtain imputations performed by dHICA.

## References

1. Brian D Strahl and C David Allis. The language of covalent histone modifications. Nature, 403(6765):41–45, 2000.

2. Tony Kouzarides. Chromatin modifications and their function. Cell, 128(4):693–705, 2007.

3. Bruce Stillman. Histone modifications: insights into their influence on gene expression. Cell, 175(1):6–9, 2018.

4. Andrew J Bannister and Tony Kouzarides. Regulation of chromatin by histone modifications. Cell research, 21(3):381–395, 2011.

5. ENCODE Project Consortium et al. An integrated encyclopedia of dna elements in the human genome. Nature, 489(7414):57, 2012.

6. Jill E Moore, Michael J Purcaro, Henry E Pratt, Charles B Epstein, Noam Shoresh, Jessika Adrian, Trupti Kawli, Carrie A Davis, Alexander Dobin, et al. Expanded encyclopaedias of dna elements in the human and mouse genomes. Nature, 583(7818):699–710, 2020.

7. Anshul Kundaje, Wouter Meuleman, Jason Ernst, Misha Bilenky, Angela Yen, Alireza Heravi-Moussavi, Pouya Kheradpour, Zhizhuo Zhang, Jianrong Wang, Michael J Ziller, et al. Integrative analysis of 111 reference human epigenomes. Nature, 518(7539):317–330, 2015.

8. Jason Ernst and Manolis Kellis. Large-scale imputation of epigenomic datasets for systematic annotation of diverse human tissues. Nature biotechnology, 33(4):364–376, 2015.

9. Timothy J Durham, Maxwell W Libbrecht, J Jeffry Howbert, Jeff Bilmes, and William Stafford Noble. Predictd parallel epigenomics data imputation with cloud-based tensor decomposition. Nature communications, 9(1):1402, 2018.

10. Jacob Schreiber, Timothy Durham, Jeffrey Bilmes, and William Stafford Noble. Avocado: a multi-scale deep tensor factorization method learns a latent representation of the human epigenome. Genome biology, 21:1–18, 2020.

11. Jian Zhou and Olga G Troyanskaya. Predicting effects of noncoding variants with deep learning–based sequence model. Nature methods, 12(10):931–934, 2015.

12. Qijin Yin, Mengmeng Wu, Qiao Liu, Hairong Lv, and Rui Jiang. Deephistone: a deep learning approach to predicting histone modifications. BMC genomics, 20:11–23, 2019.

13. Yan Li, Lijun Quan, Yiting Zhou, Yelu Jiang, Kailong Li, Tingfang Wu, and Qiang Lyu. Identifying modifications on dna-bound histones with joint deep learning of multiple binding sites in dna sequence. Bioinformatics, 38(17):4070–4077, 2022.

14. Dipankar Ranjan Baisya and Stefano Lonardi. Prediction of histone post-translational modifications using deep learning. Bioinformatics, 36(24):5610–5617, 2020.

15. David R Kelley. Cross-species regulatory sequence activity prediction. PLoS computational biology, 16(7):e1008050, 2020.

16. Žiga Avsec, Vikram Agarwal, Daniel Visentin, Joseph R Ledsam, Agnieszka Grabska-Barwinska, Kyle R Taylor, Yannis Assael, John Jumper, Pushmeet Kohli, and David R Kelley. Effective gene expression prediction from sequence by integrating long-range interactions. Nature methods, 18(10):1196–1203, 2021.

17. Alexander Karollus, Thomas Mauermeier, and Julien Gagneur. Current sequence-based models capture gene expression determinants in promoters but mostly ignore distal enhancers. Genome Biology, 24(1):56, 2023.

18. Zhong Wang, Alexandra G Chivu, Lauren A Choate, Edward J Rice, Donald C Miller, Tinyi Chu, Shao-Pei Chou, Nicole B Kingsley, Jessica L Petersen, Carrie J Finno, et al. Prediction of histone post-translational modification patterns based on nascent transcription data. Nature genetics, 54(3):295–305, 2022.

19. Zhenhao Zhang, Fan Feng, Yiyang Qiu, and Jie Liu. A generalizable framework to comprehensively predict epigenome, chromatin organization, and transcriptome. Nucleic Acids Research, 51(12):5931–5947, 2023.

20. Jason Ernst and Manolis Kellis. Chromatin-state discovery and genome annotation with chromhmm. Nature protocols, 12(12):2478–2492, 2017.

21. A Vaswani, N Shazeer, N Parmar, J Uszkoreit, L Jones, AN Gomez, L-Kaiser, and I Polosukhin. Attention is all you need in advances in neural information processing systems, 2017. Search PubMed, pages 5998–6008.

22. EA Feingold, PJ Good, MS Guyer, S Kamholz, L Liefer, K Wetterstrand, FS Collins, TR Gingeras, D Kampa, EA Sekinger, et al. The encode (encyclopedia of dna elements) project. Science, 306(5696):636–640, 2004.

23. Haley M Amemiya, Anshul Kundaje, and Alan P Boyle. The encode blacklist: identification of problematic regions of the genome. Scientific reports, 9(1):9354, 2019.

24. Lucille Lopez-Delisle, Leily Rabbani, Joachim Wolff, Vivek Bhardwaj, Rolf Backofen, Björn Grüning, Fidel Ramírez, and Thomas Manke. pygenometracks: reproducible plots for multivariate genomic datasets. Bioinformatics, 37(3):422–423, 2021.

25. Benjamin L Kidder, Gangqing Hu, and Keji Zhao. Chip-seq: technical considerations for obtaining high-quality data. Nature immunology, 12(10):918–922, 2011.

26. Xiangshuo Kong, Guisheng Wei, Nan Chen, Shudi Zhao, Yunwang Shen, Jianjia Zhang, Yang Li, Xiaoqun Zeng, and Xiaofeng Wu. Dynamic chromatin accessibility profiling reveals changes in host genome organization in response to baculovirus infection. PLoS Pathogens, 16(6):e1008633, 2020.

27. Jacob Schreiber, Carles Boix, Jin wook Lee, Hongyang Li, Yuanfang Guan, Chun-Chieh Chang, Jen-Chien Chang, Alex Hawkins-Hooker, Bernhard Schölkopf, Gabriele Schweikert, et al. The encode imputation challenge: a critical assessment of methods for cross-cell type imputation of epigenomic profiles. Genome biology, 24(1):79, 2023.

28. Chao Cheng, Koon-Kiu Yan, Kevin Y Yip, Joel Rozowsky, Roger Alexander, Chong Shou, and Mark Gerstein. A statistical framework for modeling gene expression using chromatin features and application to modencode datasets. Genome biology, 12:1–18, 2011.

29. Arjan van der Velde, Kaili Fan, Junko Tsuji, Jill E Moore, Michael J Purcaro, Henry E Pratt, and Zhiping Weng. Annotation of chromatin states in 66 complete mouse epigenomes during development. Communications Biology, 4(1):239, 2021.

30. Jacob Schreiber, Ritambhara Singh, Jeffrey Bilmes, and William Stafford Noble. A pitfall for machine learning methods aiming to predict across cell types. Genome biology, 21:1–6, 2020.

31. Yizhong Wang, Yang Li, Cankun Wang, Chan-Wang Jerry Lio, Qin Ma, and Bingqiang Liu. Cemig: prediction of the cis-regulatory motif using the de bruijn graph from atac-seq. Briefings in Bioinformatics, 25(1):bbad505, 2024.

32. Sarah Kim-Hellmuth, François Aguet, Meritxell Oliva, Manuel Muñoz-Aguirre, Silva Kasela, Valentin Wucher, Stephane E Castel, Andrew R Hamel, Ana Viñuela, Amy L Roberts, et al. Cell type–specific genetic regulation of gene expression across human tissues. Science, 369(6509):eaaz8528, 2020.

33. Shushan Toneyan, Ziqi Tang, and Peter K Koo. Evaluating deep learning for predicting epigenomic profiles. Nature machine intelligence, 4(12):1088–1100, 2022.

34. Manolis Kellis, Barbara Wold, Michael P Snyder, Bradley E Bernstein, Anshul Kundaje, Georgi K Marinov, Lucas D Ward, Ewan Birney, Gregory E Crawford, Job Dekker, et al. Defining functional dna elements in the human genome. Proceedings of the National Academy of Sciences, 111(17):6131–6138, 2014.

35. Ritambhara Singh, Jack Lanchantin, Gabriel Robins, and Yanjun Qi. Deepchrome: deep-learning for predicting gene expression from histone modifications. Bioinformatics, 32(17):i639–i648, 2016.

36. Dohoon Lee, Jeewon Yang, and Sun Kim. Learning the histone codes with large genomic windows and three-dimensional chromatin interactions using transformer. Nature Communications, 13(1):6678, 2022.

37. Arshdeep Sekhon, Ritambhara Singh, and Yanjun Qi. Deepdiff: Deep-learning for predicting differential gene expression from histone modifications. Bioinformatics, 34(17):i891–i900, 2018.

38. Jason Ernst, Pouya Kheradpour, Tarjei S Mikkelsen, Noam Shoresh, Lucas D Ward, Charles B Epstein, Xiaolan Zhang, Li Wang, Robbyn Issner, Michael Coyne, et al. Mapping and analysis of chromatin state dynamics in nine human cell types. Nature, 473(7345):43–49, 2011.

39. Matthew T Maurano, Richard Humbert, Eric Rynes, Robert E Thurman, Eric Haugen, Hao Wang, Alex P Reynolds, Richard Sandstrom, Hongzhu Qu, Jennifer Brody, et al. Systematic localization of common disease-associated variation in regulatory dna. Science, 337(6099):1190–1195, 2012.

40. Jason Ernst and Manolis Kellis. Chromhmm: automating chromatin-state discovery and characterization. Nature methods, 9(3):215–216, 2012.

41. Helga Thorvaldsdóttir, James T Robinson, and Jill P Mesirov. Integrative genomics viewer (igv): high-performance genomics data visualization and exploration. Briefings in bioinformatics, 14(2):178–192, 2013.

42. Maya Kasowski, Sofia Kyriazopoulou-Panagiotopoulou, Fabian Grubert, Judith B Zaugg, Anshul Kundaje, Yuling Liu, Alan P Boyle, Qiangfeng Cliff Zhang, Fouad Zakharia, Damek V Spacek, et al. Extensive variation in chromatin states across humans. Science, 342(6159):750–752, 2013.

43. Mehdi Foroozandeh Shahraki, Marjan Farahbod, and Maxwell W Libbrecht. Robust chromatin state annotation. Genome Research, 2024.

44. Peter M Visscher, Naomi R Wray, Qian Zhang, Pamela Sklar, Mark I McCarthy, Matthew A Brown, and Jian Yang. 10 years of gwas discovery: biology, function, and translation. The American Journal of Human Genetics, 101(1):5–22, 2017.

45. Melina Claussnitzer, Judy H Cho, Rory Collins, Nancy J Cox, Emmanouil T Dermitzakis, Matthew E Hurles, Sekar Kathiresan, Eimear E Kenny, Cecilia M Lindgren, Daniel G MacArthur, et al. A brief history of human disease genetics. Nature, 577(7789):179–189, 2020.

46. Annalisa Buniello, Jacqueline A L MacArthur, Maria Cerezo, Laura W Harris, James Hayhurst, Cinzia Malangone, Aoife McMahon, Joannella Morales, Edward Mountjoy, Elliot Sollis, et al. The nhgri-ebi gwas catalog of published genome-wide association studies, targeted arrays and summary statistics 2019. Nucleic acids research, 47(D1):D1005–D1012, 2019.

47. Yu Gyoung Tak and Peggy J Farnham. Making sense of gwas: using epigenomics and genome engineering to understand the functional relevance of snps in non-coding regions of the human genome. Epigenetics & chromatin, 8:1–18, 2015.

48. Lucas D Ward and Manolis Kellis. Interpreting noncoding genetic variation in complex traits and human disease. Nature biotechnology, 30(11):1095–1106, 2012.

49. Ricardo Cruz-Herrera del Rosario, Jeremie Poschmann, Sigrid Laure Rouam, Eileen Png, Chiea Chuen Khor, Martin Lloyd Hibberd, and Shyam Prabhakar. Sensitive detection of chromatin-altering polymorphisms reveals autoimmune disease mechanisms. Nature methods, 12(5):458–464, 2015.

50. Lei Hou, Xushen Xiong, Yongjin Park, Carles Boix, Benjamin James, Na Sun, Liang He, Aman Patel, Zhizhuo Zhang, Benoit Molinie, et al. Multitissue h3k27ac profiling of gtex samples links epigenomic variation to disease. Nature Genetics, 55(10):1665–1676, 2023.

51. Wilson Lek Wen Tan, Chukwuemeka George Anene-Nzelu, Eleanor Wong, Chang Jie Mick Lee, Hui San Tan, Sze Jing Tang, Arnaud Perrin, Kan Xing Wu, Wenhao Zheng, Robert John Ashburn, et al. Epigenomes of human hearts reveal new genetic variants relevant for cardiac disease and phenotype. Circulation Research, 127(6):761–777, 2020.

52. Richard A Gibbs, Eric Boerwinkle, Harsha Doddapaneni, Yi Han, Viktoriya Korchina, Christie Kovar, Sandra Lee, Donna Muzny, Jeffrey G Reid, Yiming Zhu, et al. A global reference for human genetic variation. Nature, 526(7571):68–74, 2015.

53. Qiao Liu, Fei Xia, Qijin Yin, and Rui Jiang. Chromatin accessibility prediction via a hybrid deep convolutional neural network. Bioinformatics, 34(5):732–738, 2018.

54. Wenjie Sun, Jeremie Poschmann, Ricardo Cruz-Herrera Del Rosario, Neelroop N Parikshak, Hajira Shreen Hajan, Vibhor Kumar, Ramalakshmi Ramasamy, T Grant Belgard, Bavani Elanggovan, Chloe Chung Yi Wong, et al. Histone acetylome-wide association study of autism spectrum disorder. Cell, 167(5):1385–1397, 2016.

55. Fabian Grubert, Judith B Zaugg, Maya Kasowski, Oana Ursu, Damek V Spacek, Alicia R Martin, Peyton Greenside, Rohith Srivas, Doug H Phanstiel, Aleksandra Pekowska, et al. Genetic control of chromatin states in humans involves local and distal chromosomal interactions. Cell, 162(5):1051–1065, 2015.

56. Mingxiang Teng and Rafael A Irizarry. Accounting for gccontent bias reduces systematic errors and batch effects in chip-seq data. Genome research, 27(11):1930–1938, 2017.

57. Pang Wei Koh, Emma Pierson, and Anshul Kundaje. Denoising genome-wide histone chip-seq with convolutional neural networks. Bioinformatics, 33(14):i225–i233, 2017.

58. Xiang Zhou, Carolyn E Cain, Marsha Myrthil, Noah Lewellen, Katelyn Michelini, Emily R Davenport, Matthew Stephens, Jonathan K Pritchard, and Yoav Gilad. Epigenetic modifications are associated with inter-species gene expression variation in primates. Genome biology, 15:1–19, 2014.

