## Supplementary for "Discriminative histone imputation using chromatin accessibility"

**Text S1. Description om evaluating imputation performance in HM Peaks**

We start by downloading ENCODE narrow peak regions for various histones across cell lines. For a specific cell line, we divide the genome into 1000-bp segments and keep those with over 50% overlap with HM narrow peaks. The retained segments are then deduplicated and saved in a bed file format, serving as the dataset to evaluate imputations, focusing on regions with at least one histone modification.

**Text S2. Description on distinguishing cell-specific HM peaks**

In determining cell-specific peaks, the genome is initially partitioned into 1000-bp windows. These are then overlapped with narrow peak regions from both K562 and GM12878 cell lines to classify them as shared or cell-specific. An interval overlapping exclusively with K562 and not GM12878 is tagged as K562-specific, and vice versa for GM12878-specific peaks.

**Text S3. Metrics for assessing chromatin state segmentation**

Chromatin state annotations are essential for many genomic tasks, including identifying active regulatory elements and interpreting disease-associated genetic variation, and it’s a main application for imputations of histone modifications, for instance, ChromHMM is a tool commonly utilized for these purposes in the ENCODE project. However, despite the widespread applications of ChromHMM and other methods, no principled approach exists to evaluate the statistical significance of their assignments.

Our evaluation leverages SAGAconf to gauge the segmentation accuracy provided by ChromHMM using dHICA and EPCOT imputations. We infer the quality of these imputations based on segmentation performance, employing three key metrics from SAGAconf, as enumerated in Table S1.

Table S1. The metrics we used.

| Name | Explanation |
| --- | --- |
| auSMC per chromatin state | The area under this state merging curve (auSMC) is a measure of a state's reproducibility when taking into account such merges, calculated according the ratio of the observed area under the curve |
| Average r-value genome-wide | This r-value is assigned to each genomic bin of a SAGA annotation and represents the probability that the label of this bin will be reproduced in a replicated experiment. |
| posterior calibration curve r-square | r-square score represents how well the isotonic regression curve fits the data. |

Table S2. Chromatin accessibility data source of experiments

| **ENCODE accession** | **Genome assembly** | **Cell line** | **Sequencing** | **Output type** |
| --- | --- | --- | --- | --- |
| ENCFF451IKJ | GRCh38 | K562 | ATAC-seq | fold change over control |
| ENCFF156WZT | GRCh38 | K562 | ATAC-seq | fold change over control |
| ENCFF093IIW | GRCh38 | K562 | ATAC-seq | fold change over control |
| ENCFF092ESJ | GRCh38 | K562 | ATAC-seq | fold change over control |
| ENCFF943HUS | GRCh38 | K562 | DNase-seq | fold change over control |
| ENCFF564YLJ | GRCh38 | K562 | DNase-seq | read-depth normalized signal |
| ENCFF352SET | GRCh38 | K562 | DNase-seq | read-depth normalized signal |
| ENCFF137BQX | GRCh38 | K562 | DNase-seq | read-depth normalized signal |
| ENCFF316PVI | GRCh38 | GM12878 | ATAC-seq | fold change over control |
| ENCFF725NBM | GRCh38 | HCT116 | ATAC-seq | fold change over control |
| ENCFF782BVX | GRCh38 | MCF-7 | ATAC-seq | fold change over control |
| ENCFF343VRK | GRCh38 | CD4 | ATAC-seq | fold change over control |
| ENCFF420BRB | GRCh38 | GM12878 | DNase-seq | read-depth normalized signal |
| ENCFF169PCK | GRCh38 | HCT116 | DNase-seq | read-depth normalized signal |
| ENCFF799DOV | GRCh38 | MCF-7 | DNase-seq | read-depth normalized signal |
| ENCFF194NJI | GRCh38 | HELA | DNase-seq | read-depth normalized signal |
| ENCSR050BJO | GRCm38 | Hindbrain | DNase-seq | read-depth normalized signal |
| ENCFF672DJH | GRCm38 | ES | DNase-seq | read-depth normalized signal |

Table S3. Mouse’ s ChIP-seq data source of experiments

| **ENCODE accession** | **HM markers** | **Cell line** |
| --- | --- | --- |
| ENCFF300TRP | [H3K27ac](https://www.encodeproject.org/targets/H3K27ac-mouse/) | Hindbrain |
| ENCFF315URY | [H3K4me1](https://www.encodeproject.org/targets/H3K4me1-mouse/) | Hindbrain |
| ENCFF637YEQ | [H3K9me3](https://www.encodeproject.org/targets/H3K9me3-mouse/) | Hindbrain |
| ENCFF437NAJ | [H3K4me3](https://www.encodeproject.org/targets/H3K4me3-mouse/) | Hindbrain |
| ENCFF058HXC | [H3K36me3](https://www.encodeproject.org/targets/H3K36me3-mouse/) | Hindbrain |
| ENCFF369FIZ | [H3K27me3](https://www.encodeproject.org/targets/H3K27me3-mouse/) | Hindbrain |
| ENCFF286QJO | H3K9me3 | ES |
| ENCFF506IKC | H3K36me3 | ES |
| ENCFF224VJA | H3K4me1 | ES |
| ENCFF955FPW | H3K4me3 | ES |
| ENCFF676VTC | H3K27ac | ES |
| ENCFF676NQE | H3K9ac | ES |

Fig. S1. The Pearson correlations of HMs across cell lines.

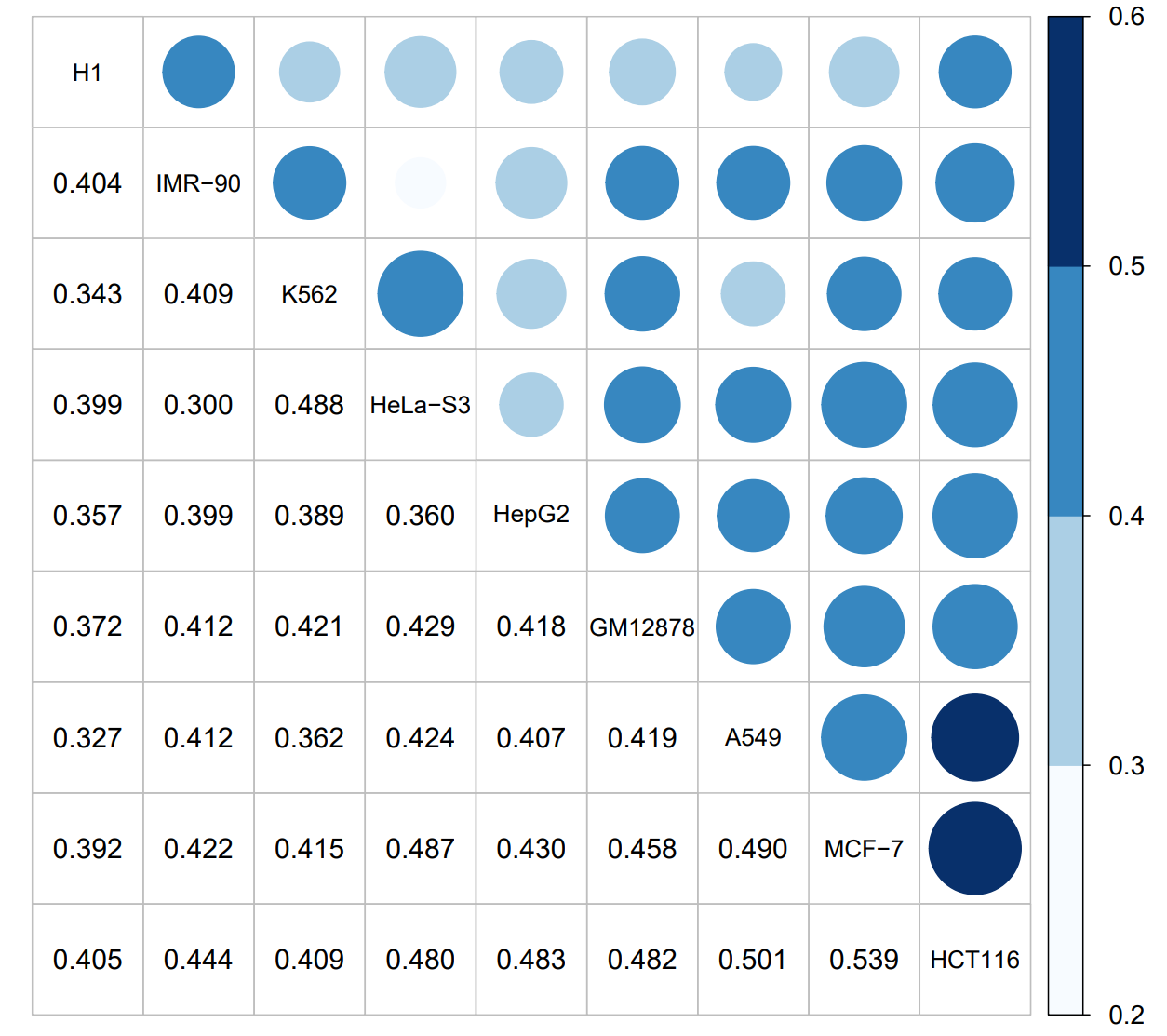

Fig. S2. Comparison of Pearson’s correlation of DNase models across cell lines, tissues and species. Empty cells indicate that no experimental data are available for comparison in the cell type shown

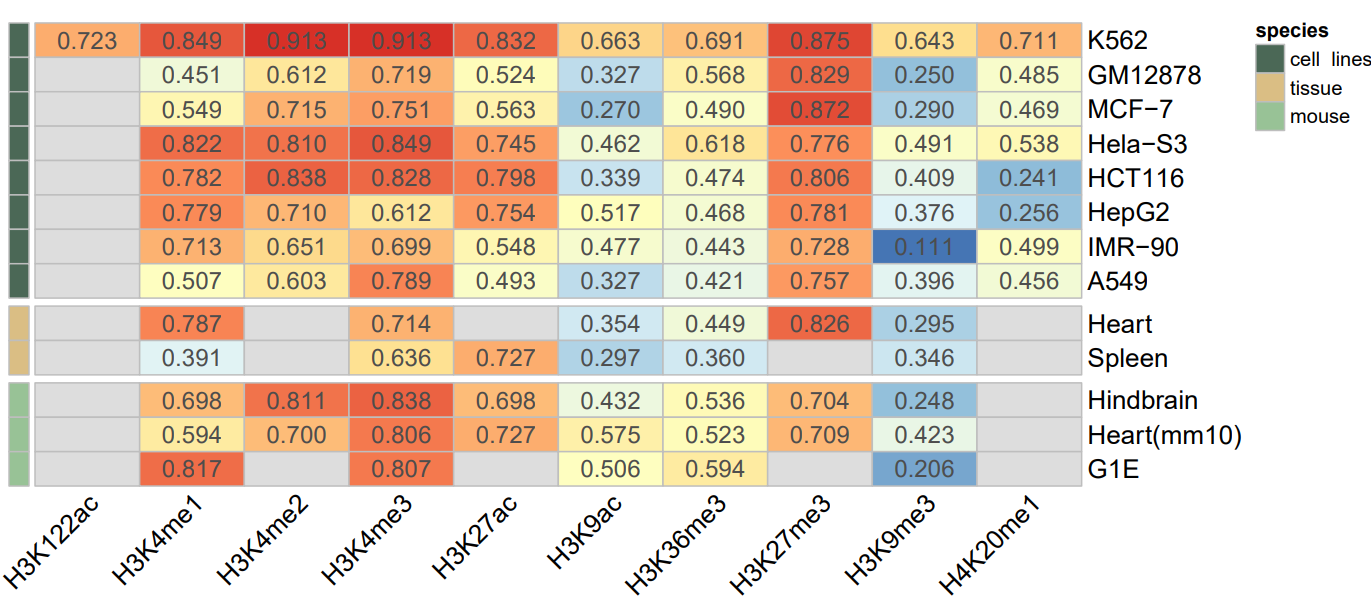

Fig. S3. The AUPRC of dHICA and the other models on K562 cell line across HM peaks.

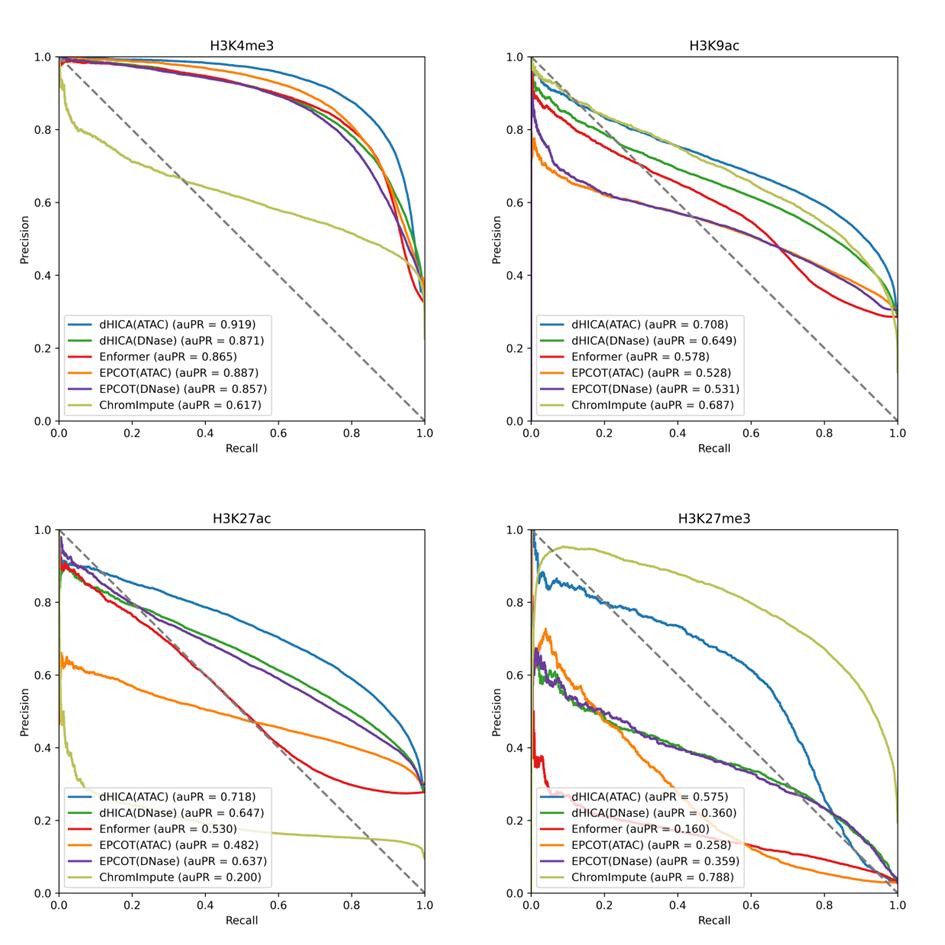

Fig. S4. The AUROC of dHICA and the other models on K562 cell line across HM peaks.

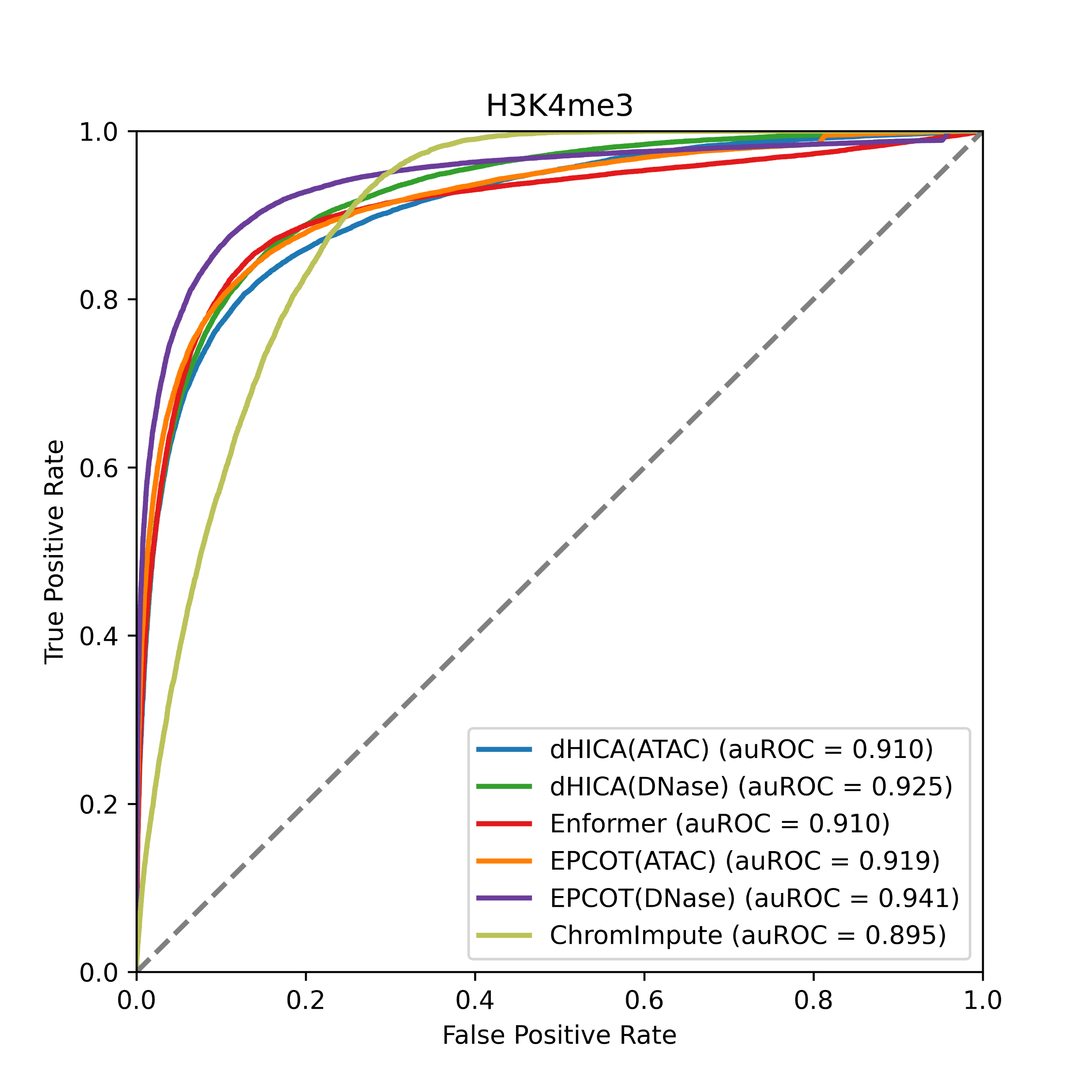

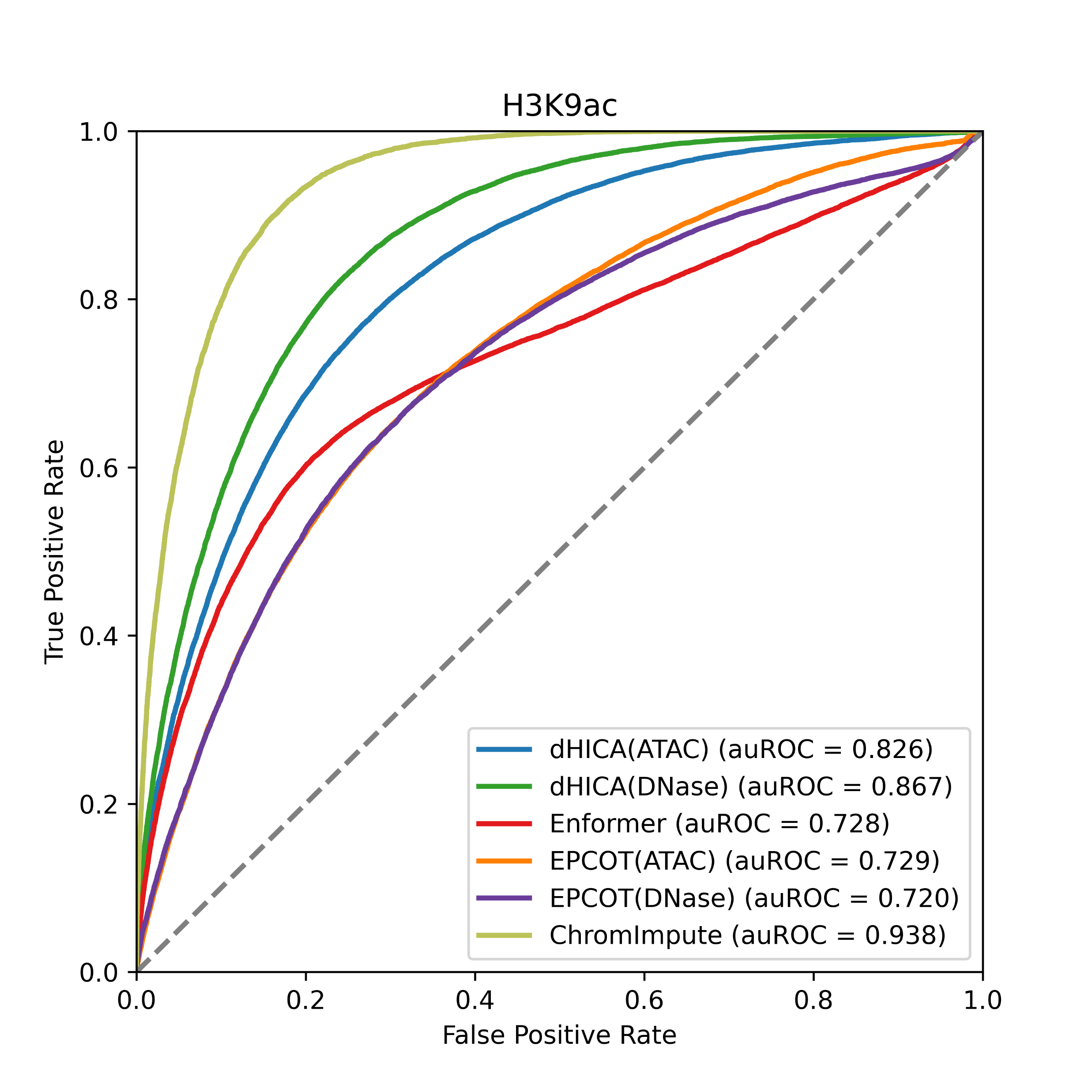

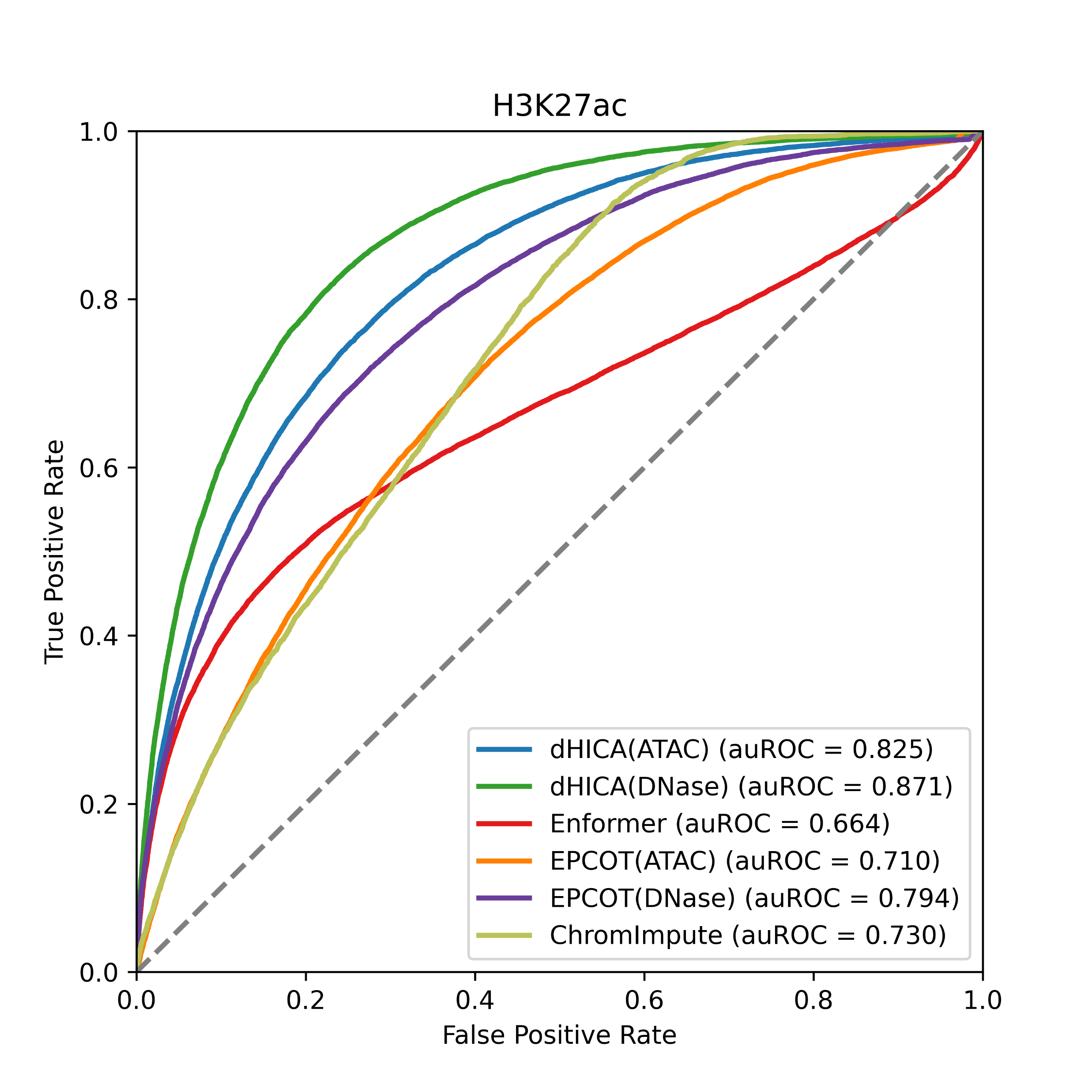

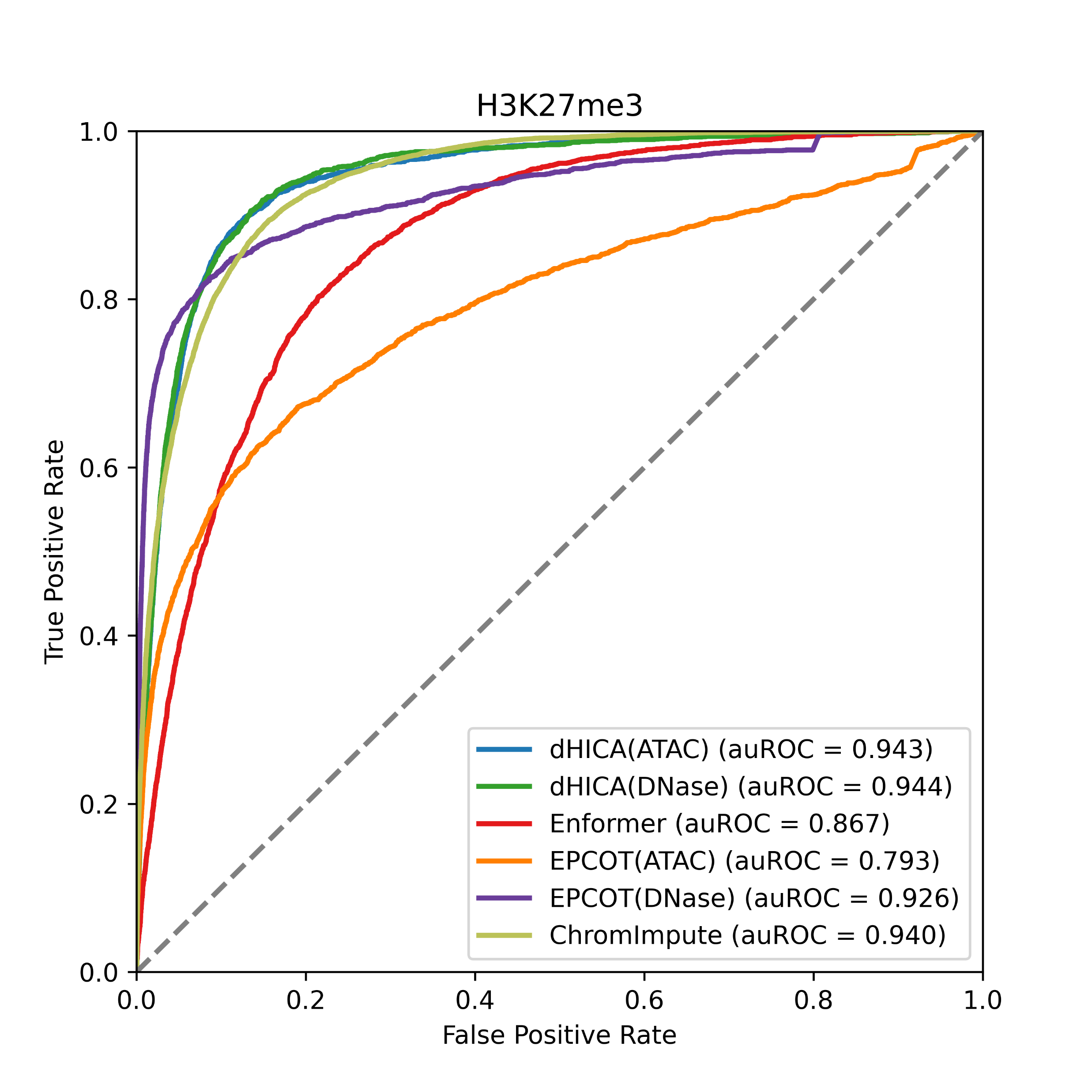
